# Humans Discriminate Individual Large-Billed Crows by their Calls

**DOI:** 10.1101/2020.09.22.308163

**Authors:** Sabrina Schalz, Thomas E. Dickins

**Affiliations:** Department of Psychology, Middlesex University, London, UK

**Author notes:** corresponding author: Sabrina Schalz.

**Keywords:** Acoustic communication, Crow call, Human–animal interaction, Individual discrimination

## Abstract

Previous research has shown that humans can discriminate two individual rhesus monkeys (*Macaca mulatta*), as well as two individual zebra finches (*Taeniopygia guttata*) by their vocalizations. The discrimination of individual zebra finches largely relies on differences in pitch contour, although this is not the only relevant cue. The purpose of the present experiment was to examine whether humans can also discriminate two individual large-billed crows (*Corvus macrorhynchos*) by their calls. Discrimination was tested with a forced-choice Same-Different Paradigm. Results show a high discrimination accuracy without prior training, although the scores obtained here were lower than those in the zebra finch discrimination task. There was no significant learning trend across trials. Future studies should investigate which acoustic cues participants use for the discrimination of individual crows and expand these findings with more non-human animal vocalizations.

## 1. Introduction

Humans readily discriminate others by their voices, primarily based on differences in mean fundamental frequency (F0, perceived as pitch), as well as the mean frequency of the first formant (F1) for female voices and mean formant dispersion (difference between F4 and F5) for male voices (Baumann & Belin 2010). Perception of these acoustic cues may vary between individuals, including individual differences in sensitivity to F0 in phoneme categorization tasks (Kong & Edwards 2016).

Individual discrimination is not limited to conspecific vocalizations but can be extended to non-human vocalizations as well. Human infants and adults can discriminate two individual rhesus monkeys (*Macaca mulatta*) by their voices, although discrimination accuracy decreased with age and adult discrimination success was only slightly above chance level (Friendly et al. 2014). Human adults are also able to discriminate two individual zebra finches by their signature songs (Schalz & Dickins, in press), which follow an individualized, stereotyped pattern (Miller 1979). After the removal of pitch contour (pitch frequency pattern), the discrimination accuracy was significantly lower, although still significantly above chance level. This suggests that pitch contour is a highly relevant cue, but not the only one (Schalz & Dickins, in press). However, this perceptual ability does not apply to all heterospecific vocalizations, as humans have been found unable to discriminate between individual dogs (*Canis familiaris*) based on their barks (Molnár et al., 2006). As dogs are able to discriminate between individual conspecifics by their barks (Molnár et al., 2009), this is due to limitations of human perception rather than lack of discrimination cues in the barks themselves. Human discrimination of individual heterospecific vocalizations is therefore not a given, highlighting the need for further studies with a diverse range of species.

Large-billed crows produce individually distinct calls and can recognize non-breeding flock members based on these calls. This signature voice system includes various F0 parameters, such as mean frequency and the slope between the start, peak and end frequency, as discrimination cues (Kondo et al. 2010). The present experiment is based on our previous discrimination experiments with zebra finch songs (Schalz & Dickins, in press) and examines whether humans can discriminate two individual large-billed crows by their individual calls.

## 2. Methods

### 2.1 Subjects

Participants (N=50, 32 female, 17 male) were students and staff at Middlesex University between the ages 18 to 50 who did not report hearing problems and gave informed consent. First-year Psychology students received credit points for participation (N=29).

### 2.2 Stimuli

Stimuli were nine calls recorded from each of two large-billed crows (both female, 4 years old). Calls were recorded by S.S. at Keio University, Tokyo. All acoustic measurements were made in Praat version 6.0.49 (Boersma & Weenink 2019; see table 1 for acoustic parameters, and figures 1 and 2 for sample spectrograms).

**Table 1:**
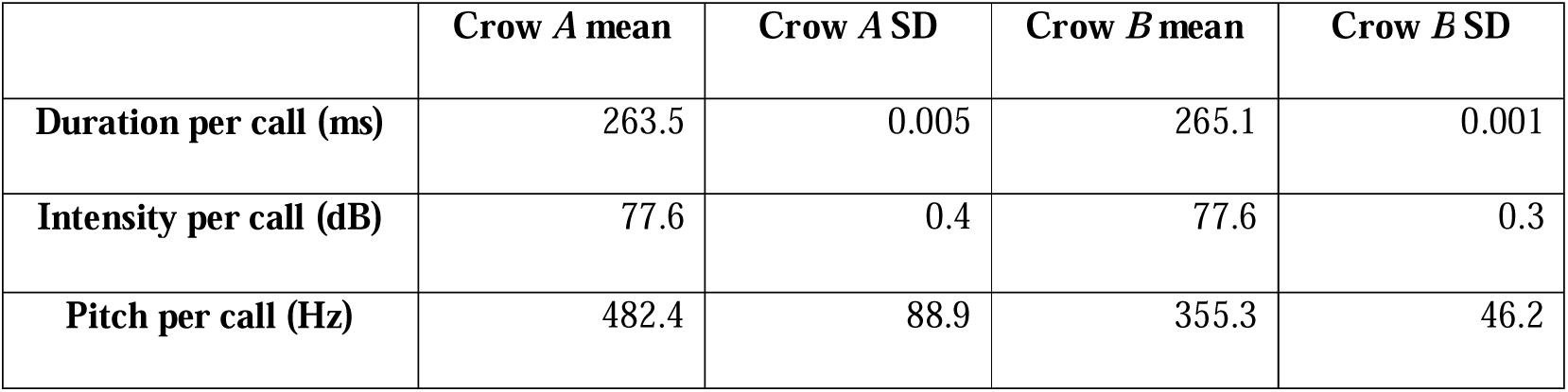
Acoustic parameters of the nine selected calls of each crow. Frequency range was set at 50Hz to 1,000Hz for pitch.

**Figure 1:**
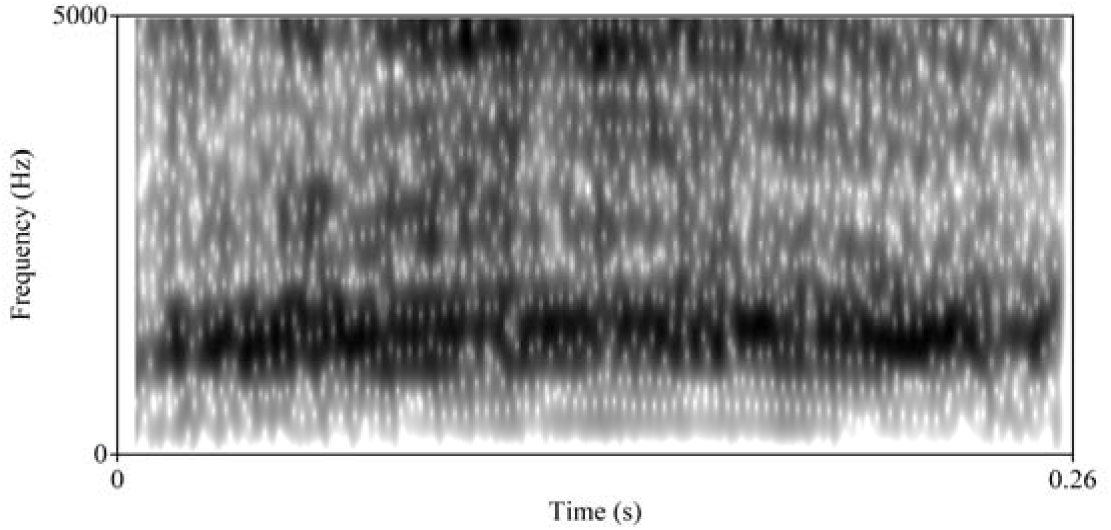
Spectrogram of sample call from crow *A*, created in Praat version 6.0.49 (Boersma & Weenink, 2019).

**Figure 2:**
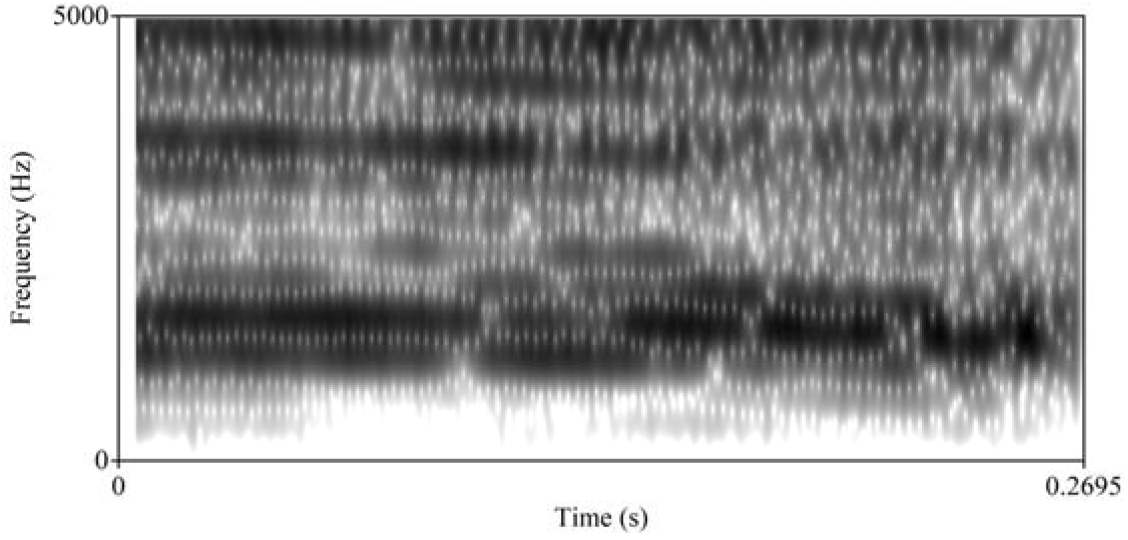
Spectrogram of sample call from crow *B*, created in Praat version 6.0.49 (Boersma & Weenink, 2019).

### 2.3 Apparatus

The experimental task and participant background questionnaire were presented in PsychoPy version 3.2 (Peirce et al. 2019) on a desktop computer in a quiet room. Stimuli were played through over-ear headphones.

### 2.4 Experimental Procedures

Discrimination accuracy was assessed with a forced-choice Same-Different Paradigm (Pisoni and Lazarus 1974) with a total of 40 trials (as in Friendly et al. 2014). Each trial consisted of two calls, either produced by the same individual (“same”-trial) or by two different individuals (“different”-trial). Pairs and trials were randomly assigned by PsychoPy. After listening to both calls, participants were asked whether they thought the call was produced by the same individual, to which they could respond “yes” and “no”. Participants did not receive training before or feedback during the experiment.

Participants’ responses were divided into four response types: “Hit” (yes on a “same”-trial), “miss” (no on a “same”-trial), “correct reject” (no on a “different”-trial), and “false alarm” (yes on a “different”-trial). The proportions of “hit” responses out of “same”-trials, and “false alarm” responses out of “different”-trials were used to calculate the discrimination sensitivity index d’ (Stanislaw & Todorov 1999) in R version 3.6.1 (R Core Team 2020), using the R package psyphy (Knoblauch 2014). The resulting d’ scores indicate the discrimination accuracy on a continuous scale from 0 (chance level, no discrimination) to 5.94 (perfect discrimination). D’ scores are an appropriate measurement for this experiment because they are less susceptible to participants’ response biases than other methods, such as the percentage of correct responses per participant (Stanislaw & Todorov 1999). D’ scores cannot be calculated with absolute values for hit rates and false alarm rates, so three “hit” rates and five “false alarm” rates were corrected following formula 1 as described in (Snodgrass & Corwin 1988).

1. 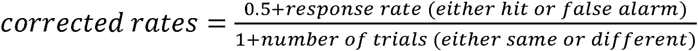

### 2.5 Statistical Analyses

Whether d’ scores were significantly above chance level was tested with a one-sample t-test (mu=0). D’ scores were also compared with those obtained in the previous experiment with zebra finch songs (Schalz & Dickins, in press) using a Mann-Whitney U test. Whether discrimination accuracy improved over the course of the trials was examined with a Mann-Kendall trend test with the R package “Kendall” (McLeod, 2011). Data for this analysis consisted of the percentage of correct responses (either “hit” or “correct reject”) for a given trial pooled from all participants.

## 3. Results

The average d’ score was 2.48 (SD=1.1, 95% CI [2.17, 2.78]), and d’ scores were significantly above chance level (t=15.86, df=49, p<0.01). Compared with the d’ scores reported for the discrimination of zebra finches (Schalz & Dickins, in press), d’ scores obtained in the present experiment were significantly below those obtained with natural zebra finch song (W=669, d=0.9, p<0.01), and significantly above those obtained for song without pitch contour (W=1024.5, p<0.01, d=1.22, p<0.01; see figure 3).

**Figure 3:**
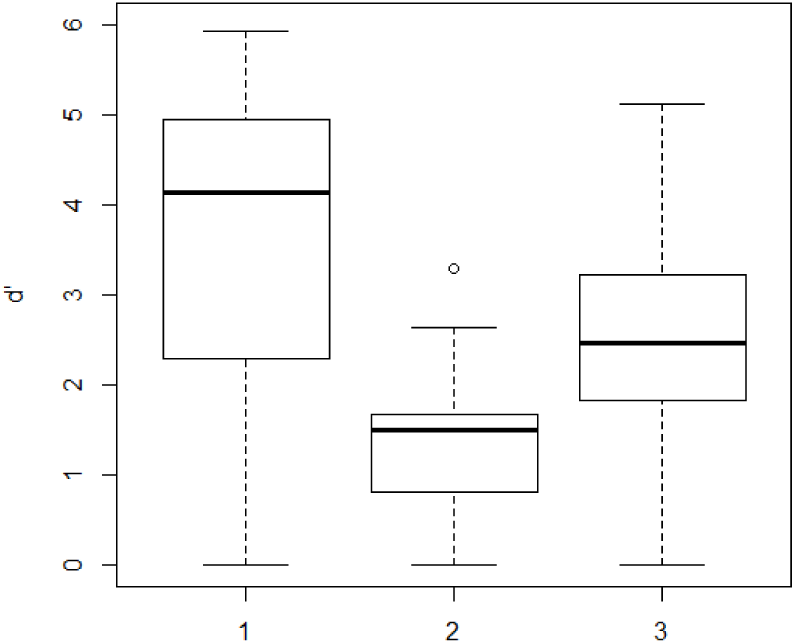
d’ scores compared between natural zebra finch songs (1), zebra finch songs without pitch contour (2; Schalz & Dickins, in press), and crow calls (3). Mean d’ scores are 3.68, 1.3, and 2.48 respectively.

The Mann Kendall Trend test showed a non-significant, although positive trend across all trials (τ =0.19; see figure 4).

**Figure 4.**
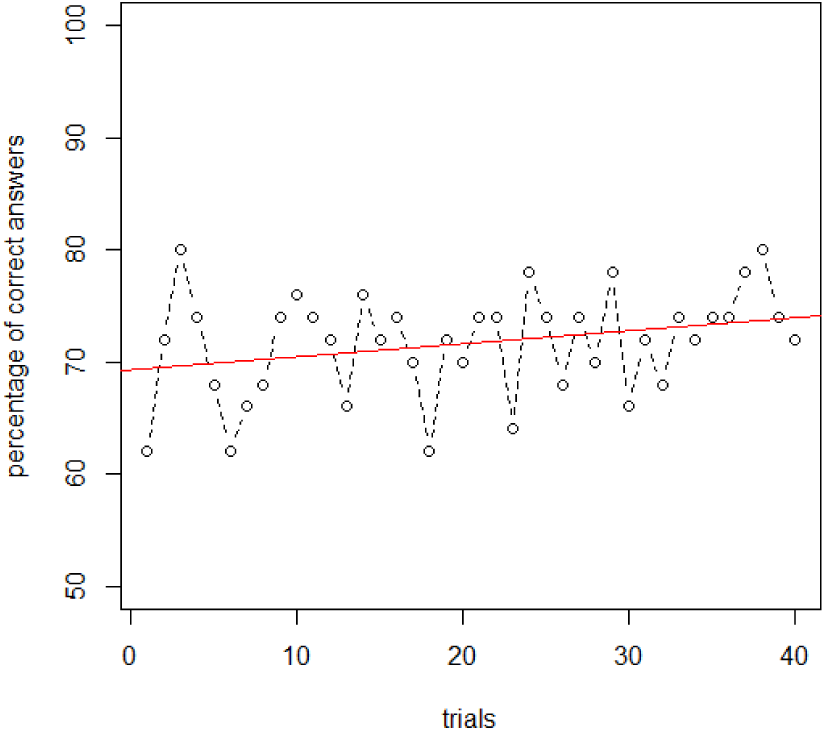
Percentage of correct answers (either “hit” or “correct reject”) for each trial pooled from all participants per trial, with a linear regression line indicating overall trend.

## 4. Discussion

These results show that humans can discriminate individual large-billed crows by their calls, although there was considerable variation in discrimination accuracy between participants (see figure 3). The mean d’ score reported here (2.48) is significantly below the one reported for the individual discrimination of zebra finches (3.68) but significantly higher than the ones for zebra finches without pitch contour (1.3, see figure 3; Schalz & Dickins, in press). This may be due to the more stereotyped pattern in natural zebra finch songs, which likely facilitates the discrimination but is lost after pitch contour removal. It remains to be investigated which acoustic cues humans use to discriminate individual crows. The mean d’ score for crow calls is also considerably higher than the reported d’ score for rhesus monkey vocalizations (0.37; Friendly et al. 2014). While this discrepancy may in part be due to methodological differences, it is nevertheless a considerable difference in discrimination accuracy.

As with the zebra finch discrimination, there was no significant learning trend across trials and there seems to be no meaningful improvement of discrimination accuracy, which was already relatively high in the first trials. It is however conceivable that accuracy would eventually improve after additional practice. It is also possible that some participants’ individual accuracy improved over the course of these trials, which would not necessarily appear in the pooled performance analysed here. Due to the small number of calls sampled from only two individuals, we cannot exclude the possibility that these two crows by chance have unusually distinct calls, and that the discrimination could be more difficult with other individuals of this species. This experiment nevertheless provides a valuable proof of concept and shows that human heterospecific acoustic discrimination is worth further investigation. To date the number of heterospecific vocalizations studied in this context is very small, and it remains unclear how discrimination accuracy and the relevance of available discrimination cues vary between the vocalizations of different species. It is noteworthy that the two experiments with heterospecific mammal vocalizations found that human adults struggle with individual discrimination (Friendly et al. 2014; Molnár et al., 2006), whereas the two experiments with avian vocalizations show the contrary. More work is needed to establish whether this is can be generalized to all mammals and birds and if so, whether individual discrimination is exceptionally high for bird vocalizations due to the similarities between birdsong and human speech (Doupe & Kuhl 1999).

## Acknowledgements

We thank Ei-Ichi Izawa for enabling the incidental recording of the calls during an unrelated project (Schalz & Izawa, 2020) that was carried out as part of the “Thesis@Keio” programme at Keio University and financially supported by JSPS KAKENHI #17H02653 and Keio University Grant-in-Aid for Innovative Collaborative Research Projects #MKJ1905 to E.-I. I.

## Declarations

### Ethics approval

The experiment was approved by the Middlesex University Psychology Research Ethics Committee (application number 8360).

### Consent to participate

All participants gave informed consent prior to the experiment.

### Consent for publication

All participants gave informed consent prior to the experiment.

### Availability of data and material

All data generated or analysed during this study can be found here: https://doi.org/10.6084/m9.figshare.12746105.v1

